# Density-Dependent Cell Migration in the Absence of Social Interactions: A Case Study of *Acanthamoeba castellanii*

**DOI:** 10.1101/2025.03.16.643508

**Authors:** Nasser Ghazi, Mete Demircigil, Olivier Cochet-Escartin, Amandine Chauviat, Sabine Favre-Bonté, Christophe Anjard, Jean-Paul Rieu

## Abstract

Cell migration is often influenced by intercellular or social interactions, ranging from long-range diffusive cues to direct contacts that can trigger biochemical signaling within the cell and affect the cell protruding activity or direction of turns. Here we study the density-dependent migration of the amoeba *Acanthamoeba castellanii (Ac)*, a unicellular eukaryote that moves without social interactions. Using experiments and mean free path theory, we characterize how collisions affect motility parameters in crowded environments. We identify the collision rate as a key parameter linking cell density to the collision-induced reorientation rate, and we show its consistency across multiple independent approaches. Our findings reveal that the intrinsic migration speed remains constant, while persistence time and effective diffusion are entirely governed by collisions. At high densities, cells exhibit nearly ballistic trajectories between collisions, a behavior rarely reported in eukaryotes. These results establish *Ac* as a minimal model for motility in the absence of biochemical signaling, with implications for testing behaviors in complex crowded environments and pre-jamming dynamics.

## Introduction

Cell migration is a fundamental process observed in living systems. In animals, it is a cornerstone of numerous biological processes, including immune responses (1), cancer metastasis, and tissue morphogenesis (2). Cell migration is also crucial for bacteria and protozoa (amoeba, protists) as it enables survival and adaptation; in bacteria, it facilitates processes like biofilm formation and infection (3), while in protozoa, it supports environmental lifecycle transitions (4), navigation, and predation (5). Cells within organisms, microbes in biofilms, soil, or the gut are not isolated; they constantly interact with each other and with the generally porous or fibrous structure of the microenvironment. Social interactions between cells largely affect individual random cell motility (speed, diffusion coefficient, or persistence time) but as well chemo- or other taxis contributing to the emergence of collective behaviors.

There are two broad classes of mechanisms through which cells detect the presence of others and modulate the magnitude and orientation of their movement in response:

The first class of mechanism is long-range sensing by diffusive cues, *i*.*e*. chemical factors that are secreted by some cells and detected by nearby cells. Quorum sensing factors (QSF) are such secreted molecules that accumulate in the cell surroundings at a rate proportional to cell density. In bacterial systems, a QSF concentration above a given threshold modifies gene expression, single cell behavior and eventually the whole population functions (6). Cancer cells may also use QSF to regulate multicellular functions and control steps in metastatic colonization (7). In MDA-MB-231 breast cancer cell lines, interleukin IL-6/8 signaling amplified by cell density does cause faster migration (8). The social amoeba *Dictyostelium discoideum* (*Dd*) is a convenient model system to study how social interactions and in particular secreted factors regulate motility and development. Upon starvation, *Dd* cells start to secrete cAMP (cyclic adenosine monophosphate), which functions as an extracellular regulatory molecule binding to cell-surface receptors of neighboring cells that activate intracellular signal transduction pathways. The cAMP relay system controls cell streaming and aggregation in *Dd*, cellular differentiation (9, 10) as well as the expression of a number of classes of genes (9, 10). Some of us found that in its vegetative phase (*i*.*e*., in presence of food), *Dd* motility is downregulated by a QSF complex larger than 60 kDa (11, 12).

In the second class of mechanisms, physical contact between cells allows for short-range sensing of neighbors either triggering - not mutually exclusively - cell-cell adhesion, cell-cell repulsion, cell repolarization, or just excluded volume (EV) interactions. EV refers to the steric effects that arise when two or more physical objects cannot occupy the same space simultaneously. In the context of cell migration on two-dimensional (2D) surfaces, this means that a cell’s movement is constrained by the physical presence of other cells provided that cells do not overlap in the direction normal to the surface. EV interactions are purely repulsive and generate avoidance between cells but do not necessarily lead to active re-routing of their trajectories or significant shape deformation after the cells in contact move away (13).

By contrast, we will use the terms “collisions” or “contact interactions” interchangeably to describe contact events leading to active reorientations of cell trajectories due to active cell shape changes and protruding activity. Among the various types of contact interactions, contact inhibition of locomotion (CIL), where cell contact redirects movement away from the collision position has been observed in various cellular models, including chick fibroblasts and neural crest cells (14, 15). CIL was not observed in *Dd*, but some of us proposed a variant called contact enhancement of locomotion (CEL), where cell collisions momentarily enhance the persistence of cells (13). Finally, when cells express some cell-cell adhesion molecules, they tend to follow others in streams or small cohorts and to align (16, 17).

Earlier studies at cellular high packing fractions with slowly moving cells (epithelial or mesenchymal cells) have reported contrasted results: cell speed and persistence can decrease with increasing density as in nearly confluent epithelia (18), stay the same for 3T3 fibroblast cells (19) or even increase for breast cancer cells (20). Understanding the nature of the different interactions at play in these experiments is challenging because interactions usually combine. Numerical simulations from the active matter community could be very useful to test hypotheses. Some of us have widely used active Brownian particle (ABP) (13) or agent-based models (21) to interpret complex experiments with various intermingled interactions.

To address these problems of intricate interactions, we propose *Acanthamoeba castellanii* (*Ac*) as a simpler model organism. In contrast to the social amoebae like *Dd* forming multicellular aggregates and fruiting with spores that can remain dormant in the absence of nutrients (4), *Ac* forms solitary cysts that allow survival in nutrient poor conditions (22). In a comparative study between *Dd* and *Ac*, it was shown that *Ac* movement is primarily determined by neither chemotaxis nor cell-cell interactions (23). *Ac* hence represents a strikingly different paradigm as a solitary amoeba, maintaining solitary behavior even in dense environments. This distinction offers a unique opportunity to explore the effects of cell density and contact interactions on migration in a system where chemical signaling is minimal, and cells are not adhering to each other. The life cycle of the *Acanthamoeba* consists of two stages, a trophozoite stage in favorable conditions where cells present an active amoeboid motility and a cyst stage under adverse environmental conditions where cells become double-walled, overall round but with a slightly wrinkled outer cyst wall and a polyhedral inner cyst wall (22). Relatively little is known about the movement of *Ac* (23, 24).

In this work, we study how cell density modulates *Ac* migration in the absence of long-range signaling. In high-density environments, *Ac* cells experience frequent collisions that modulate their trajectories, leading to a nearly ideal density-dependent migration characterized by nonlinear scaling of mean-squared-displacements parameters with density. To the best of our knowledge, this is the first experimental study showing clearly the influence of collisions on cell diffusion via the mean free path model.

## Materials and methods

### Cell culture and cell line

For our study, we used the axenic free-living strain Acanthamoeba castellanii (Ac), ATCC NEF 30010 isolated from soil in the United States; California; Pacific Grove. They were grown in proteose peptone-yeast-glucose (PYG90) medium (25) as monolayers in 75 cm^2^ tissue culture flasks at 25°C. Prior to the experiment, adherent cells were harvested and resuspended to a stock concentration of 4 × 10^6^ amoeba per milliliter.

### Videomicroscopy

Cell behavior was monitored using three distinct microscopes. At low cell densities, where cells would readily leave the FOV in other microscopes, a “lens-free” Cytonote 6W microscope (IPRASENSE, Montpellier), which can simultaneously image six stationary samples over a wide field of approximately 30 mm^2^ at a resolution of 1.68 µm/pixel was used. The Cytonote employs an integrated holographic reconstruction algorithm (HORUS software) that automatically preprocesses the images, eliminating the need for manual focusing. A TE2000-E inverted microscope (Nikon) controlled with Micro-Manager (version 2.0 gamma) and equipped with a motorized stage, a 4× Plan Fluor objective (Nikon), and a Zyla camera (Andor) was used for bright field timelapse imaging at high densities (*ρ* > 600cells/mm^2^). Moreover, for cell shape analysis, we took images in transmission mode with a confocal microscope (Leica SP5) operated with LAS software (version 3.4, Leica) and equipped with a 10× objective lens for higher-magnification experiments. In all three setups, images were collected at intervals compatible with cell motility timescales (5 to 60 s/frame) and, once acquired, were segmented using a machine-learning-based approach implemented in Ilastik (26). This workflow allowed us to reliably extract both cell outlines (for cell shape and collision analysis) and cell centroids (for tracking and motility analysis) across varying cell densities, image resolutions and fields of view.

### Tracking and MSD calculation with filtering criteria

To quantify single-cell trajectories in our random motility assays, we used the open-source Python library *Trackpy* (27) and standard scientific libraries *NumPy* (28), *pandas* (29), *matplotlib* (30) and *SciPy* (31). Briefly, for each frame in the recorded timelaspe series, we identified cell coordinates via the “Find Maxima” function in *ImageJ* (version 1.53q) (32) and then linked these positions into trajectories across consecutive frames. Linking was performed with a conservative search range (10-40 pixels) depending on the frame rate and a short-term “memory” (4 frames across all datasets), allowing for the occasional dropout of cell centroids without prematurely ending trajectories. Once the trajectories constructed, we computed each trajectory’s mean squared displacement (MSD) in two dimensions (2D) as a function of the lag time *t* using overlapping intervals (33) and finally the average of single cell MSD over all cells as ⟨*MSD*(*t*)⟩ =⟨(*x*_*i*_(*t*_0_−*t*)−*x*_*i*_(*t*_0_))+ (*y*_*i*_(*t*_0_−*t*)−*y*_*i*_(*t*_0_))^2^⟩, where the angle brackets ⟨ ⟩ denote the ensemble average over all cells and all starting times *t*_*o*_. We further applied a “*β*-filter” in log-log space to exclude highly super-diffusive or sub-diffusive behavior (*e*.*g*., floating or stuck cells). Specifically, we fit each individual cellular MSD curve to a power law ≈ *t*^*β*^ and kept only those with *β*_*min*_ ≤ *β* ≤ *β*_*max*_. We chose a *β*_*min*_ = 0.25 and a *β*_*max*_ = 1.75 to provide a good balance between retaining genuinely motile cells and removing outliers. To extract diffusion coefficients (*D*), we fit the ensemble MSD curves to Fürth’s model (Eq. 1) using standard non-linear least squares, *SciPy*’s curve fit (31). All other fits in this work were also done with *SciPy*’s curve fit, with uncertainties and 95% confidence intervals for parameters obtained by propagating the standard errors from the covariance matrices via the delta method (34). The velocity autocorrelation function (VACF) was computed by taking dot products of velocity vectors at different time lags and normalizing (35). To obtain the persistence time, the normalized VACF was then fitted to a single-exponential decay of the form exp(-t/τ_P_). The characteristic decay constant τ_*p*_ was interpreted as the persistence time.

### Cell-cell collisions: detection and rate analysis

We conducted collision analyses in motility experiments spanning 6 hours, each performed on a 1 mm^2^ region with an additional 50 µm buffer zone to ensure a homogeneous distribution of *Ac* cells. During these recordings, any two cells within a distance threshold equal to twice their radius, 2r, were deemed to be colliding. To efficiently detect and count all such collision events over time, we employed the *cKDTree* algorithm from the *scipy*.*spatial* module in Python (31). This method significantly reduced computational complexity by rapidly searching for nearest-neighbor pairs at each frame, thereby enabling reliable estimates of collision rates as a function of cell density.

## Results

Figure 1 and Supplementary Movie M1 illustrate the impact of cell density on the morphology and motility of *Acanthamoeba castellanii* (*Ac*) cells. At low density (30 cells/mm^2^, Fig. 1a), cells exhibit a characteristic amoeboid shape with one or two broad protrusions (pseudopods), sometimes bifurcating, retracting or just steadily expanding as cells move forward. A few rounded, and darker cells are also present at low density. These cells do not deform and either stay in place or float in the medium. They are probably cysts. The mean diameter of the *Ac* cells used in our mean free path model described later, *i*.*e*., 2*r*=30±1 μm, is obtained from such a low-density experiment. At high density (450 cells/mm^2^, Fig. 1.b), the proportion of round cells increases, suggesting a stress-induced transition to a quiescent cyst-like state. The impact of density on motility is highlighted in Figs. 1c-d and Supplementary Movie M2, which show the time-normalized trajectories over two hours. At low density, cells traverse the field of view with relatively smooth, persistent movements, gradually changing direction over time. In contrast, at high density, trajectories become erratic, with frequent zig-zag patterns and abrupt reorientations resulting from frequent collisions (red dots in Fig. 1c-d). The low-density experiment allows a clear observation of these directional changes induced by occasional contacts. These findings underscore the role of collisions in *Ac* migration, where cell-cell interactions strongly influence movement (and the number of non-moving cells), despite the absence of social signaling.

**Fig. 1.**
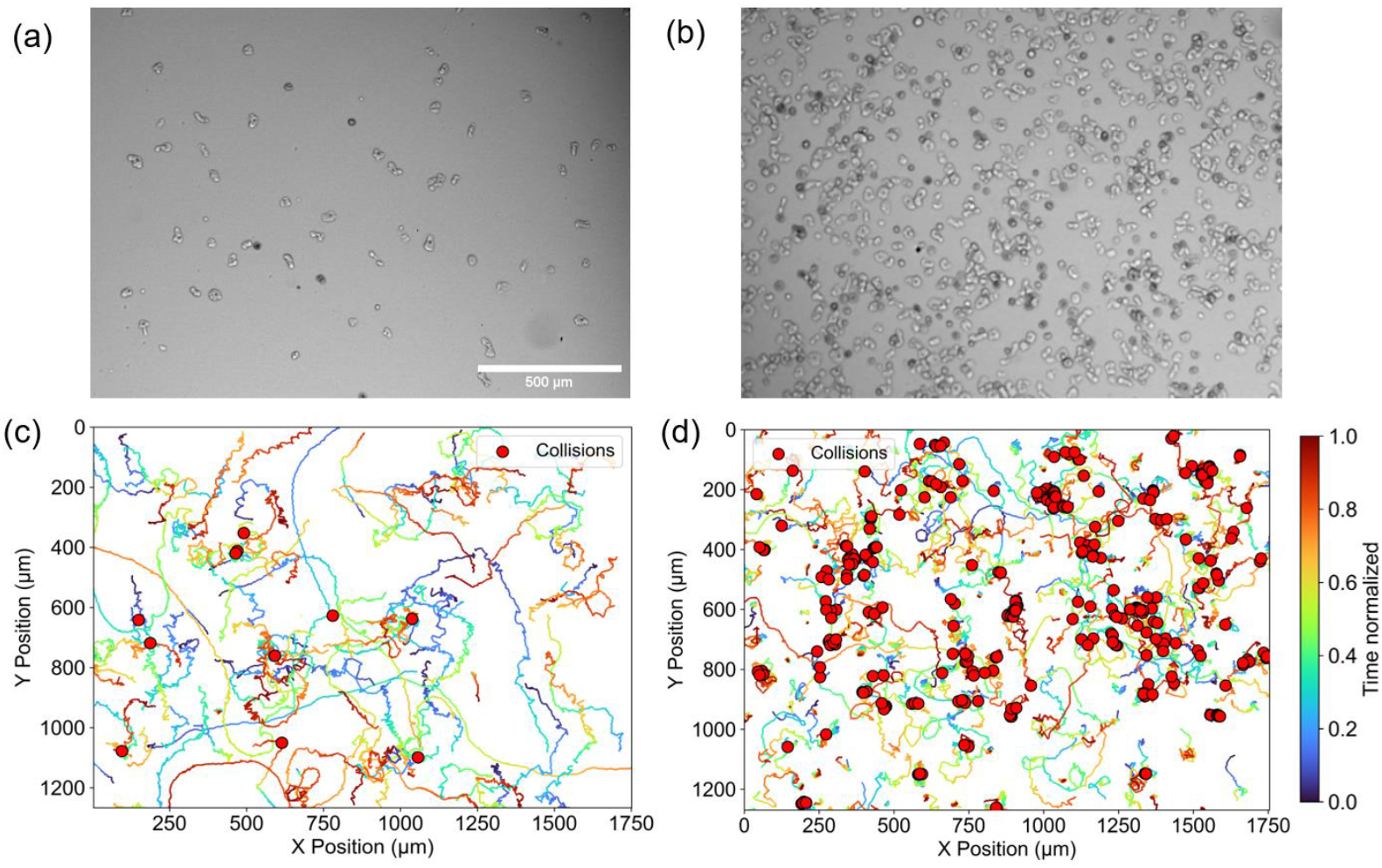
First frame images of *Ac* cells in random motility experiments taken via transmission microscopy in **a** with *ρ* = 30cells/mm^2^ and in **b** with *ρ* = 450cells/mm^2^. In **c** and **d**, the time-normalized trajectories over two-hours of **a** and **b**, with the collision events, 11 and 359 collisions, represented as red dots, respectively. Microscope: confocal microscope (Leica SP5) 10X magnification with resolution 3.03 µm/pixel.

The intricate trajectories of Fig. 1c resemble a tangled plate of spaghetti and can be quantitatively described as “correlated random walks” (CRWs). In a CRW, a particle moves persistently in a given direction for a characteristic timescale, known as the persistence time (τ), before randomly reorienting itself (36). This distinguishes CRWs from purely Brownian motion, where directional changes occur at every time step. To quantify the degree of randomness in these trajectories and characterize the transition from persistent to diffusive motion, we will compute the mean squared displacement (MSD), a key metric that captures how the average displacement of cells evolves over time. In the absence of bias, the MSD are linear at times much larger than τ with a slope corresponding to *4D* in two dimensions where *D* is the diffusion coefficient (36). To ensure reliable parameter estimation, prior work on *Dd* (11) suggested that a minimum of 200 trajectories, each lasting around 10 times τ, is required for accurate estimation of *D* (within 10% relative error) but that the time-lapse interval is only important to capture the short times. For low-density experiments, a preliminary test indicated that τ was around 30 minutes, we hence conducted 6-hour recordings at around 1 frame/min, ensuring trajectories exceeded the minimal requirement for reaching the linear diffusive regime. In order to obtain several hundred trajectories at very low cell density (ρ ≤ 10 cells/mm^2^), we used a lens-free microscope with a very large field of view (FoV) of 30 mm^2^ (Fig. 2b). This microscope records diffraction patterns of cells in 6-well plates placed directly on its 6 sensors. These patterns are then computationally reconstructed to form an in-focus image of the sample at 1.68 µm/pixel (see how this imaging technique renders single cells in Fig. 2a, which is a zoom of Fig. 2b). Besides its large FoV, this microscope is able to simultaneously compare six different densities without stage motion. Figures 2c and 2d display the reconstructed cell trajectories corresponding to the FoVs of Figs. 2a and 2b respectively. These trajectories illustrate the range of displacements observed over a two-hour period and are further visualized in Supplementary Movie M3, where individual cell motions and their directional changes can be clearly appreciated. On the other hand, for high-density experiments, we used a conventional microscope in transmission mode taking shorter time-lapse intervals to ensure proper tracking, and shorter total recording times because τ was found to be much smaller than at low density.

**Fig. 2.**
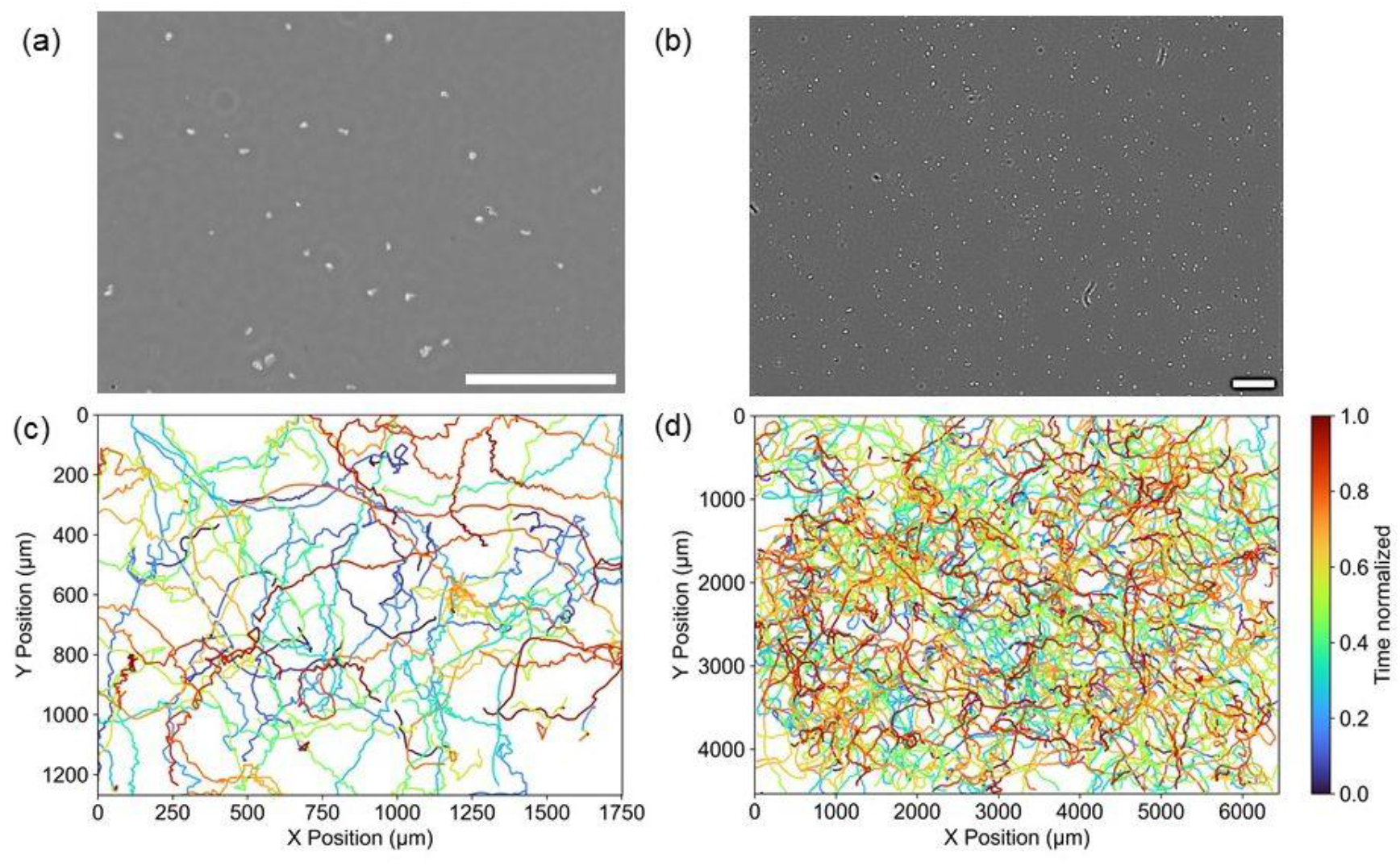
Images of Ac cells in a single random motility experiment taken via lens-free microscopy in **a** (cropped FOV) and **b** (full FOV) with *ρ* = 12cells/mm^2^. In **c** and **d**, the time-normalized trajectories over two-hours of **a** and **b**, respectively. Microscope: lens-free (Cytonote 6W) with 1.68 µm/pixel resolution and a time interval of 45s between consecutive frames. Scale bar: 500 µm.

Figure 3 presents the mean squared displacements (*MSD*) and velocity autocorrelation functions (VACF) of *Ac* at different densities. The MSD were averaged using overlapping time intervals and over all cells in a given density condition (11). Cells were discarded from this average if their MSD was too sub-diffusive or too super-diffusive (see materials and methods) in order to exclude cyst cells or floating cells. Fig. 3a-c shows the *MSD* divided by *4t* (*t* is the lag time), a convenient representation that highlights the asymptotic plateau at long times, corresponding to the diffusion coefficient *D*, and the characteristic time to reach this plateau, related to the persistence time. The *MSD* are fitted using Fürth’s model (37):

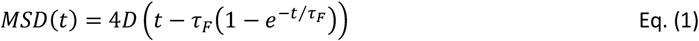

**Fig. 3.**
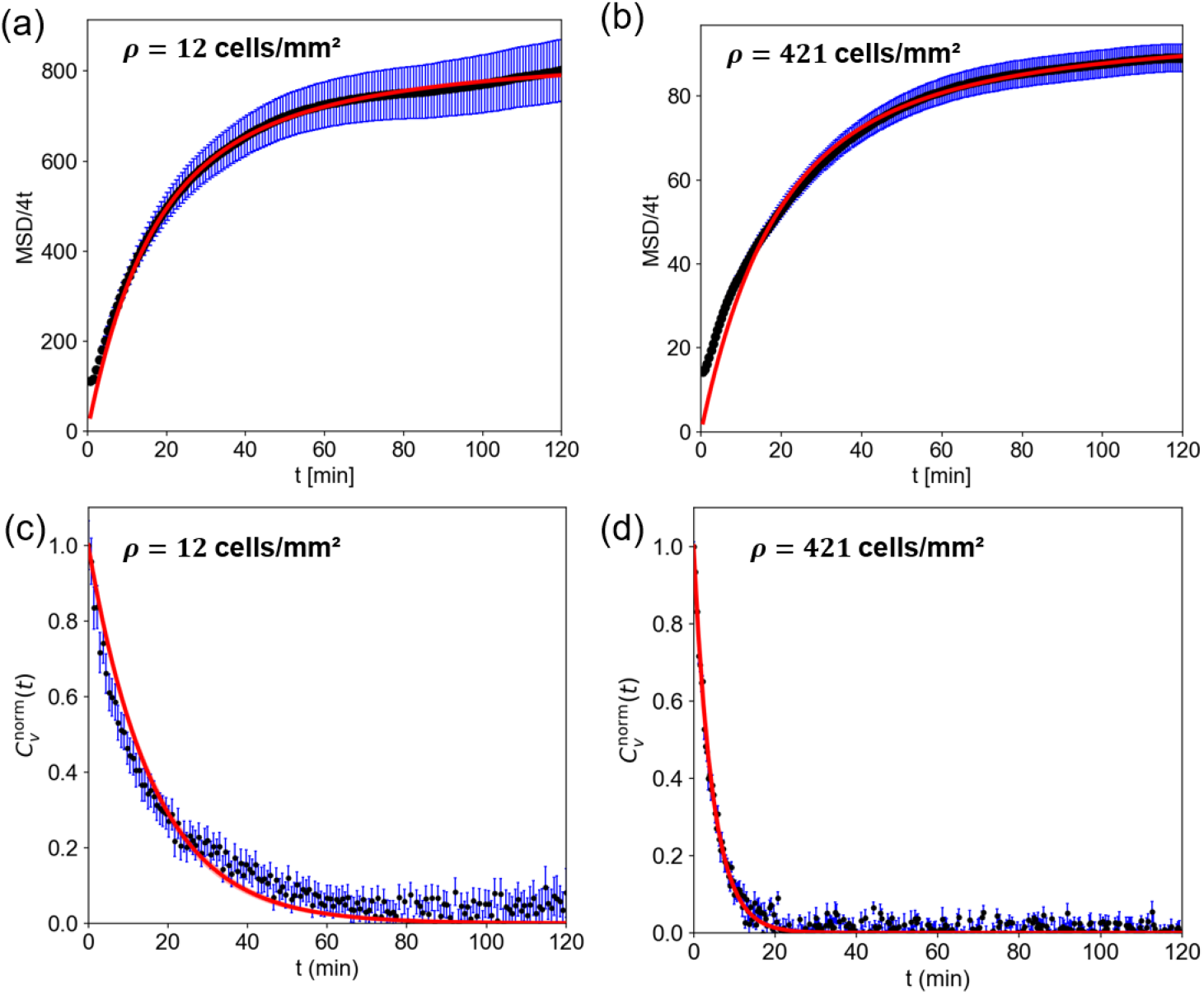
*MSD/4t* and normalized VACF of *Ac* cells in random motility experiments taken at various densities (ρ=12 and 421 cells/mm^2^ in **a, c** and **b, d** respectively). In **a-b**, the computed mean and SEM of the *MSD/4t* are in black and blue respectively. The accompanying Fürth model fits are in red: in **a**, *D*=846±3 µm^2^/min and τ_*F*_=21.9±0.2 min and in **b**, *D*=94±3 µm^2^/min and τ_*F*_=5.8±0.2 min. In **c-d**, the normalized VACF computations, 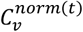, are shown in black with SEM (in blue) and exponential decay fit in red. In **c**, τ_*p*_ =16.3±0.4 min and in **d**, τ_*p*_=4.9±0.1 min. Errors on *D* and on τ_*p*_ come from 95% confidence intervals on the fits.

This equation accounts for both the ballistic regime at short times, 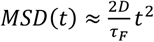, and the diffusive regime at long times, *MSD*(*t*) ≈ 4*D*(*t*−*τ*_*F*_), where τ_F_ is the Fürth’s persistence time. The extracted *D* values, *D*=846 µm^2^/min at 12 cell/mm^2^ (Fig. 3a), and *D*=93 µm^2^/min at 421 cell/mm^2^ (Fig. 3b) indicate that increasing density strongly reduces the diffusion coefficient. The Fürth’s persistence time decreases as well: τ_*F*_=21.9 and 5.8 min with increasing density respectively.

Figs. 3c-d show the normalized velocity autocorrelation functions (VACF), measuring the directional memory of cell trajectories. The decay follows an exponential law 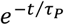 (fit in red), yielding an independent estimate of the persistence time noted here τ_p_, which also decreases with density: τ_p_ =16.3 and 4.87 min respectively (Fig. 3c-d). Note that the exponential fit of VACF is always close to experimental data while the Fürth fit systematically deviates from MSD data at short times indicating a likely short-time diffusive behavior not analyzed here (38). Overall, the VACF, which is the second derivative of the MSD, is not sensitive to such a short time regime and enables a reliable estimate of the persistence time. Fürth’s model remains valid at intermediate and long times and is thus used to extract diffusion coefficients. Increasing cell density leads to more constrained movement, shorter directional persistence, and lower effective diffusion, likely due to steric hindrance and increased cell-cell interactions.

To better quantify the effect of density on cell motility, we repeated the same experiments at intermediate densities and extracted the resulting diffusion coefficients. We observe a monotonic decrease of the diffusion coefficient with cell density. It is reduced by over 30-fold across the explored range of roughly 5 to 900 cells/mm^2^ (Fig 4a). Similarly, as cell density increases, the persistence time τ_*p*_ decreases by a factor around 30 (Fig 4b). Since the effective diffusion coefficient results from both the cells’ persistence times and their instantaneous speeds *V*_*C*_, we checked the effect of density on this parameter and find that *V*_*C*_ is also reduced by density, albeit by a smaller factor (Fig 4c).

**Fig. 4.**
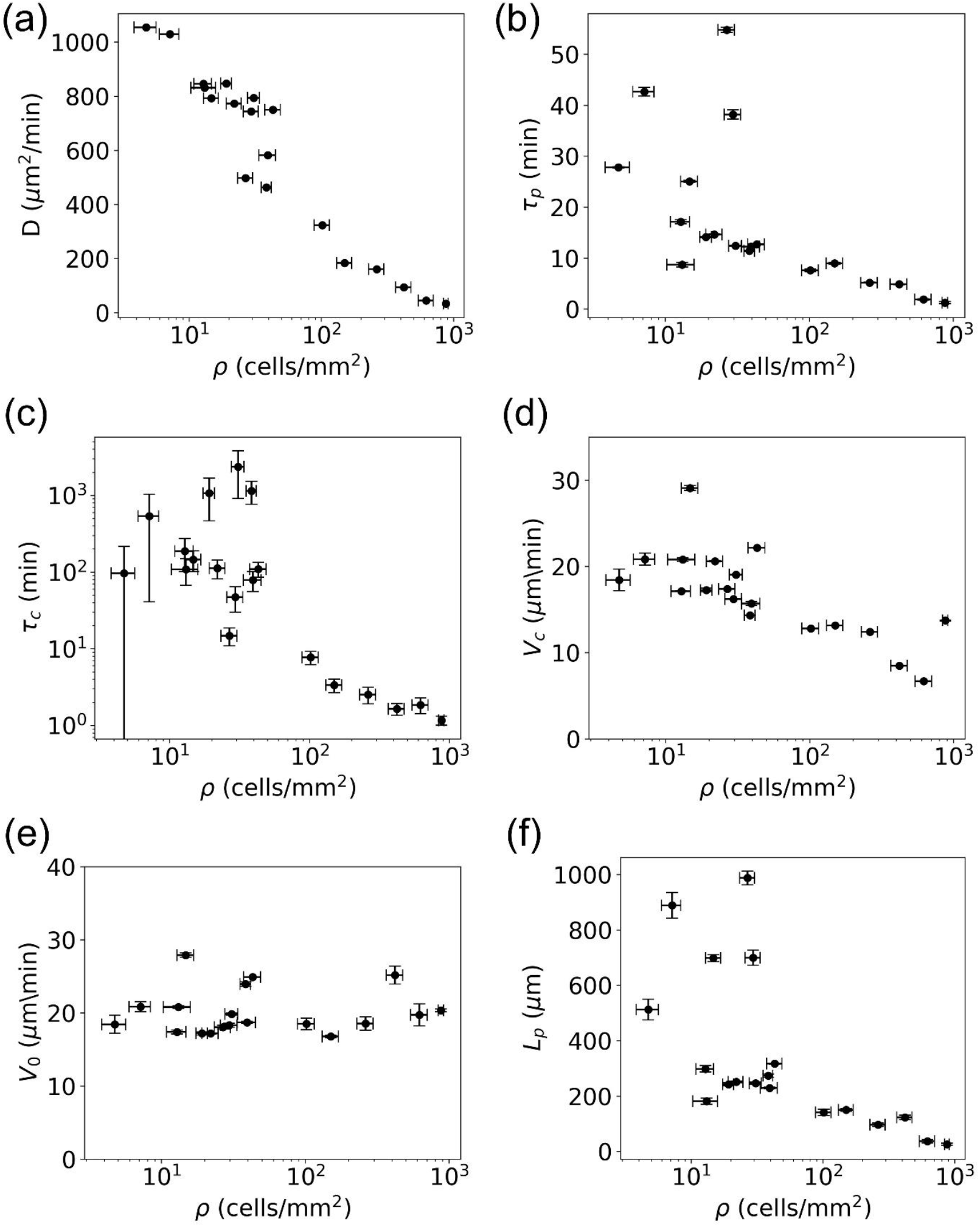
*Ac* single cell motility parameters at increasing cell densities. For all plots **a-f** circles represent the ensemble mean of the underlying parameter for a given experiment at a given cell density *ρ* and the x-axis is shown in log scale. **a**, diffusion coefficient *D* obtained by Fürth fit of MSDs. **b**, persistence time τ_*p*_ obtained from VACF exponential decay fits. **c**, mean instantaneous velocity *V*_*c*_ computed from cellular trajectories in their entirety. **d**, average time between cellular collisions τ_*c*_ obtained by directly measuring contacts between cells over time in motility experiments. Note that the y-axis is shown in log scale. **e**, mean instantaneous velocity *V*_0_ of cells for which no collisions are recorded within 5 minutes time windows. **f**, persistence length*L*_p_ = *V*_0_τ_*p*_. In **a** and **b**, the vertical error bars represent the 95% confidence intervals obtained from the Fürth (*D*) and exponential decay (τ_*p*_) fits, respectively. In **c-e**, the error bars correspond to the standard error of the mean (SEM) of the measured observables. In **f**, the error bars represent the uncertainty in the persistence length calculated via error propagation from the SEM of *V*_0_ and the 95% confidence interval of τ_*p*_.

These more refined measurements allow us to examine the origin of this dependence on density and to compare it to the predictions of simple models.

We quickly eliminated two possibilities discussed in the introduction: contact enhancement of locomotion (CEL) (13) and regulation by quorum sensing factors (QSF). To test for CEL, a thorough comparative study of cell trajectories before and after a collision in a low-density situation (ρ≤50 cell/mm^2^) showed that cell velocity and directionality (as defined by the CME, or coefficient of motion efficiency (13)) increased by less than 10% after the collision (39). In our previous studies with the social amoeba *Dd* we showed that QSF accumulate in the medium in a density-dependent manner and lower the diffusion coefficient *D* as time passes (11). Here, we used data at intermediate densities, for which we have the best statistics and calculated *D* on sliding 2h intervals during the 6h experiment. *D* presented a small increase of about 10% over 6 hours (39), possibly compatible with cumulative CEL effects, but again nothing comparable with the large variations of *D* observed in Fig. 4a.

The most obvious remaining candidate to explore was the role of collisions in the persistence time of *Ac* cells as we observe a clear reorientation after each collision (Fig. 1 and Supplementary Movies M1-M2). We started by measuring the time between collisions averaged over all cells in a specific density condition. As expected, we observe that this time is much shorter at high densities than at low ones (Fig 4d). We also notice that, at high densities, this time is only a few minutes, clearly shorter than the persistence time of *Ac* at low densities. Since cells are expected to change directions after a collision because of steric hindrance and the need to extend new protrusions, this is a strong indication that collisions are the main reason for the decrease in a cells’ persistence time. Although a cell might be in a persistent phase, it would collide with another faster than its natural persistent time, lowering it accordingly.

These collisions are so frequent that we hypothesize they bias our measurement of instantaneous speeds, as they are expected to occur between consecutive acquired frames. The resulting change in direction would therefore lead to an underestimation of the cells’ actual speed. Thanks to our automated detection of collisions, we could repeat speed measurements excluding all instances of collisions and only keep time steps where the cells were freely moving. We find that their collision-free instantaneous speed *V*_0_ is independent of density and remains constant *V*_0_ =20 ± 3 µ*m*/*min* (mean±sem, Fig 4e). Combining τp and *V*_0_, we can estimate the correlation length *L*p = *V*_0_τ_p_, *i*.*e*. the straight portion of the trajectory before a reorientation which, obviously, follows the persistence time dependence but is a convenient quantity to discuss when density effects become important (Fig. 4f).

These different results give us a simple understanding of the process: at low densities, cells move at *V*_0_ with a persistence time set by their internal migration mechanisms. When density increases, frequent collisions with neighbors prevent them from staying ballistic for extended periods, and their effective persistence time is decreased in a density-dependent way. This simple view can be quantitatively tested thanks to our accumulated experimental data. The mean free path theory indeed provides a comprehensive framework to study these effects (40).

We first explore the predictions of this theory when cells are considered purely ballistic. The mean free path *Λ* is defined as the average distance a cell travels before encountering another cell and undergoes a collision. In our two-dimensional system, it can be written as:

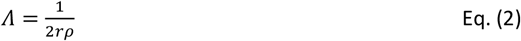

with *r* is the typical radius of one cell and the distance 2*r* corresponds to the distance between two cells interacting by contact effects (Fig 5a). This expression gives a quantitative basis to explore density-dependent effects. In our system however, the obstacles encountered by a cell, *i*.*e*. other cells, are also moving. One must therefore consider the relative velocity of cells with respect to one another. Considering two different cells with identical speeds *V*_0_ and velocities 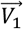 and 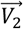, their relative velocity is 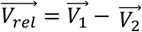 and, as 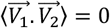 (both cellular velocities are assumed to be random and independent), the resulting RMS (root mean squared) of the relative velocity can be expressed as 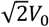. Combining the mean free path of the cell with this relative velocity allows us to extract the rate of collisions for cells at a given density as

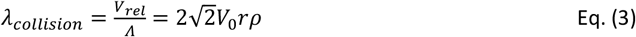

**Fig. 5.**
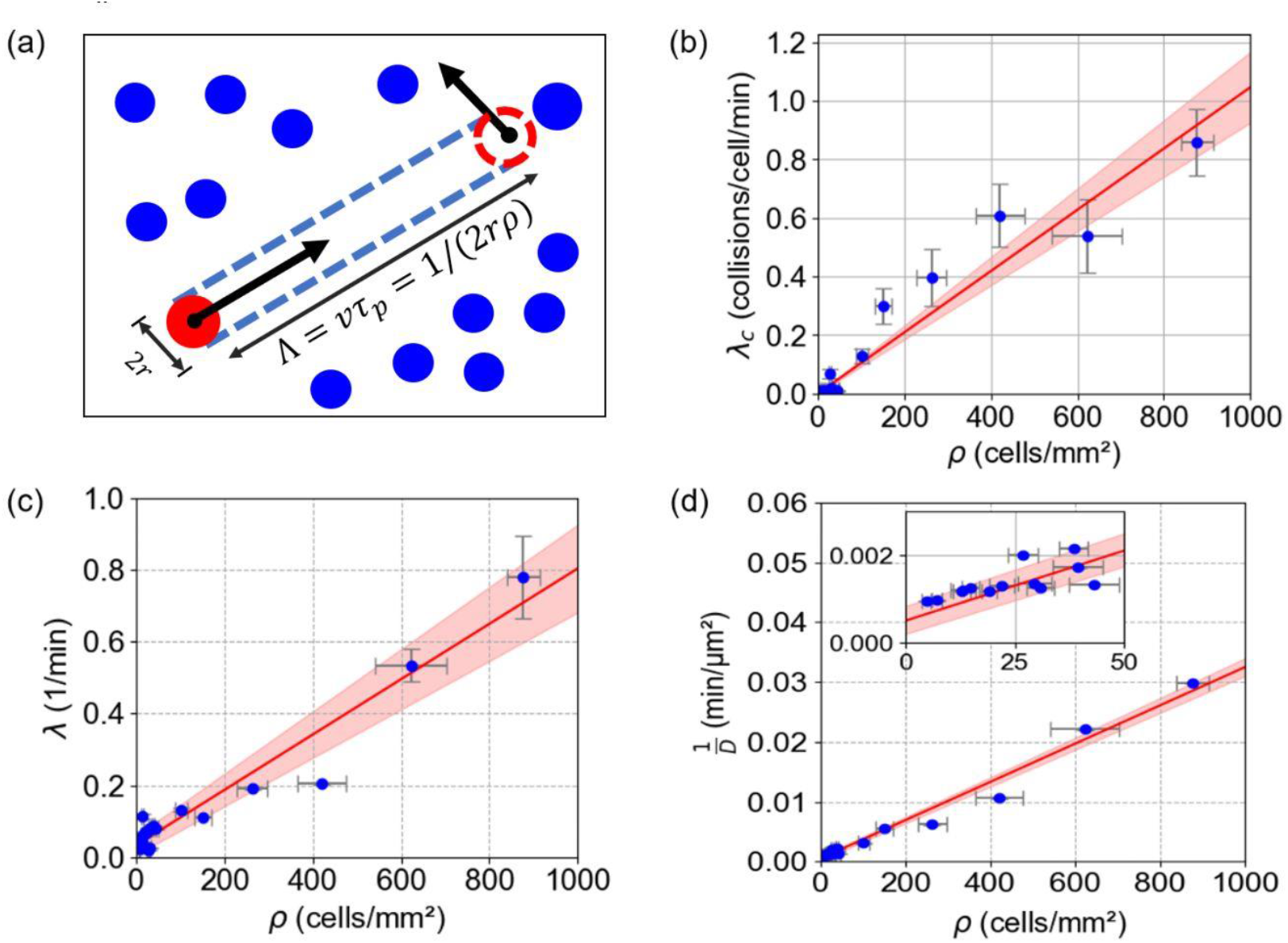
Mean free path model interpretation of the data. **(a)** Scheme representing the mean free path model relating collision events, the traveled path before a collision event Λ in a system, to the total cell density *ρ* and cell size 2*r*. **(b-d)** Plots of collision-related measured quantities as a function of cell density *ρ* for a set of 19 experiments (blue points) and linear fits (red lines) with 95% confidence intervals of the fits (light red shading). **(b)** Collision-induced reorientation rate λ_*C*_ obtained by counting the number of collisions per cell per minute (fit: λ_*C*_ = 0.00105ρ min^-1^ where *ρ* is in cells. mm^−2^) The vertical error bars represent the SEM of the number of collisions per cell per minute. **(c)** Mean total reorientation rate λ = 1/τ_*p*_ (fit: λ = 0.00077ρ + 0.0332 min^-1^ where *ρ* is in cells. mm^−2^). **(d)** Inverse of the diffusion coefficient *D* (fit: 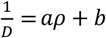 with *a* = (29 ± 1) * 10^−6^ min.mm^2^.µm^-2^.cell^-1^ and *b*=(7 ± 4)*10^−4^ min.µm^-2^ where *ρ* is in cells.mm^2^). **(b)** Vertical error bars represent the SEM of the number of collisions per cell per minute. **(c)** Vertical error bars show the 95% confidence intervals (CI) for 1/τ_*p*_. These were derived from the 95% CI of τ_*p*_ itself by 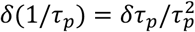. **(d)** Vertical error bars reflect error propagation from the SEM of *D* using δ(1/D) = δD/*D*^2^. In **(b-d)**, the horizontal error bars denote the SEM of cell density ρ.

As density vanishes, so does the rate of collisions. We introduce the collision rate coefficient 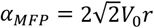 given by the mean free path. Using *V*_0_ =20 ± 3 µm/min (Fig 4e) and *r* = 15 ± 0.5μ*m*, we obtain an estimate α_*MFP*_ = (8.5 ± 1.5) * 10^−4^ collisions.mm^2^.min^-1^.cell^-2^.

To test these predictions, we look at the rate of collisions observed experimentally from the average time between cellular collisions, *i*.*e*.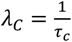, and find it to be well-fitted by a linear relationship λ_*C*_ = *α*_*C*_*ρ* (Fig. 5b). This yields a first experimental measurement of the collision rate coefficient *α*_*C*_ = (10.5 ± 1.2) * 10^−4^ collisions.mm^2^.min^-1^.cell^-2^ in good agreement with the expected value from the mean free path.

Cells, however, are not ballistic and have an intrinsic reorientation rate, which we note λ_0_ (related to their collision-free intrinsic persistence time). The total reorientation rate λ(ρ) of a cell can thus be written as:

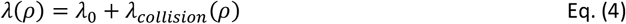

where, by definition, λ(*ρ*) is the inverse of the observed persistence time τ_*p*_(*ρ*) which can be deduced from the velocity auto-correlation. Here too, we find that its dependence in density is well fitted by a linear relationship λ(*ρ*) = λ_0_ + α_*P*_*ρ* (Fig 5c) and yields a value of α_*P*_ = (7.8 ± 1.2) * 10^−4^ collisions.mm^2^.min^-1^.cell^-2^ close to α_*C*_ obtained from collisions measurements and to *α*_*MFP*_. In addition, we obtain from the fit the intrinsic reorientation rate λ_0_ = 3.3 * 10^−2^ min^-1^ corresponding to a nearly 30 min intrinsic persistence time in the absence of collisions.

Finally, assuming the short-time asymptotic limit of the Fürth equation is verified experimentally, we expect the following relation between experimental diffusion coefficient, experimental velocity at short times and persistence 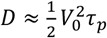. This is, however, a rough estimate as the Fürth equation does not perfectly fit data at short times (Fig. 3b-c, *i*.*e*. the MSD are not perfectly ballistic there). Adding the *ρ* dependence of τ_*p*_, we end up with the following relation where the inverse of the diffusion coefficient increases linearly with cell density:

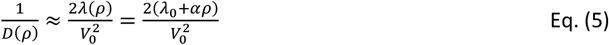

This is indeed what we observe (Fig 5d). The fit 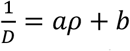 presents a slope *a* = (29 ± 1) * 10^−6^min.mm^2^.µm^-2^.cell^-1^ which is rather close to the value 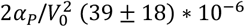 min.mm^2^.µm^-2^.cell^- 1^; the discrepancy being mostly attributed to the non-ballistic behavior of the MSD at short times. All this analysis presented in Fig. 5 strongly suggests that the effects of density are well captured by the sole effect of a reduced mean free path, with cell-cell collisions inducing changes in directions and lowering the persistence time.

To recapitulate, the good numerical agreement among the three estimates of the collision rate coefficient *α* (*α*_*MFP*_ from mean free path theory, *α*_*C*_ from direct collision counts, and *α*_*P*_ from persistence analyses) highlights the robust impact of collisions on *Ac* motility. These findings provide a strong basis for examining, in the following discussion, how steric constraints dominate the density-dependent driven migration of *Ac* cells.

## Discussion

In this study, we have demonstrated how collision dynamics govern the migration of *Ac* in the absence of social interactions. Through a comprehensive analysis employing mean free path theory and experimental observations we have shed light on the intricate ways in which collisions impact the motility of *Ac* cells at varying densities. The collision rate coefficient *α* emerged as a crucial parameter, linking cell density *ρ* to the collision-induced reorientation rate, and was consistently determined through analytical calculations from the mean free path (*α*_*MFP*_), direct experimental measurements of collisions (*α*_*C*_), and from velocity autocorrelation function analyses (*α*_*P*_). The close agreement among these obtained values of *α* helps to further validate our framework: as *Ac* cell density increases, cells collide more and more, decreasing the mean free path and influencing motility parameters. Notably, our results reveal that, despite variations in density, the intrinsic velocity *V*_0_ of individual cells in the absence of collisions remains unchanged. This invariance allows us to characterize *Acanthamoeba castellanii* motility solely in terms of collision-driven persistence. The effective diffusion coefficient *D* follows the fundamental relation 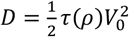 where the persistence time τ(*ρ*)is directly controlled by the collision frequency. Such a behavior is rarely observed in eukaryotic cells and establishes *Ac* as a promising model for active matter. At high densities, cell trajectories, which are linear between collisions, become erratic due to collisions. The mean free path sets the persistence length *L* = *V*_0_τ(*ρ*) ≈ *V*_0_/*αρ*, decreasing with cell density (Fig. 4f). At low density, the minimum persistence length is set by the natural persistence time τ_0_ = λ^−1^ = 30*m*i*n* and *L*_0_ = *V*_0_τ_0_ ≈20×30 = 600µ*m*. By comparison *Dictysostelium discoideum* is slower (*V*_*0*_ *≤ 7* µm/min) and less persistent (τ_0_*≤ 3* min) (13, 21). As a result, the persistence length of *Dd* is less than 20 µm (*i*.*e*., about two *Dd* cell sizes), which is 30 times smaller than that of *Ac*. Since collision effects manifest at cell densities on the order of 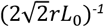 or higher, we can observe density effects as soon as *ρ* ≳ 40 cell/mm^2^ for *Ac* (*r=*15µm), whereas for *Dd* (*r*=5µm) it occurs around 3500 cell/mm^2^, corresponding to near confluence.

The implications of these findings extend beyond single-cell motility. The physics of diffusion in dense or disordered environments is a topic of broad interest in biological and active matter physics, particularly in contexts such as bacterial transport in porous media, collective escape dynamics (*i*.*e*., panic escape and crowd motion) and pre-jamming dynamics (41). The topic of bacterial diffusion in disordered and porous media is relevant to many health and environmental processes, and many recent papers tackled it. For instance, the diffusion of the swimming bacteria *Magnetococcus marinus* in micropatterned chambers with regular obstacles was measured and analyzed with a mean free path theory, similar to our approach (42). Similar analyses remain scarce and not yet detailed for eukaryotic cells (43), which are slower and far less persistent than bacteria. Future work exploring *Ac* migration under such microfabricated pillar arrays could be more conclusive in probing the effects of geometric constraints and obstacles on motility.

Additionally, an open question remains regarding the nature of post-collision scattering: does *Ac* follow a purely random reorientation, or do its trajectories exhibit characteristics akin to contact inhibition of locomotion (CIL) (15) or contact enhancement of locomotion (CEL) (13) Our analysis of velocity and CME before and after collisions (39) as well as the fact that *V*_0_ is remarkably constant regardless of density (Fig. 4e) indicates that CEL only has a minor role in the dynamics studied. Another ongoing avenue of research concerns the consequences of such a strong dependence of cell persistence on collisions for *Ac* aerotaxis, which is our main research topic (21). Preliminary experiments studying the navigation of *Ac* in self-generated or imposed oxygen gradients strongly suggest that aerotaxis also depends on cell density (39). The same study conducted on the social amoeba *Dd* did not show such a *ρ* dependence (44).

## Conclusions

This study contributes to a broader understanding of active matter dynamics in biological systems by highlighting the role of contact-mediated interactions in solitary amoebae. Insights gained from *A. castellanii* may have implications for understanding the physical constraints governing cell motility in non-cooperative systems, as well as for the principles underlying migration in high-density environments.

## Abbreviations

AC: *Acanthamoeba castellanii*
Dd: *Dictyostelium discoideum*
EV: excluded volume
2D: two-dimensional
MSD: mean squared displacements
VACF: velocity autocorrelation function

## Statements and Declarations

### Data availability

The data that support the findings of this study are available from the corresponding author upon reasonable request.

### Author contributions

N.G., C.A. and J.P.R. contributed to the study conception and design. Material preparation, data collection and analysis were performed by N.G., A.C. and M.D. The first draft of the manuscript was written by J.P.R. and O.C.E. and all authors commented on previous versions of the manuscript. All authors read and approved the final manuscript. J.P.R. supervized the work.

## Supplementary Movie captions

**Supplementary Movie M1. top** and **bottom**: two-hour movies of experiments presented in Fig. **1a** and Fig. **1b**, at *ρ* = 30cells/mm^2^ and *ρ* = 450cells/mm^2^, respectively. Scale bar: 100µm. Time stamp: hour:min

**Supplementary Movie M2. a** and **b**: two-hour movies of experiments presented in Fig. **1a** and Fig. **1b**, at *ρ* = 30cells/mm^2^ and *ρ* = 450cells/mm^2^, respectively, with ID-based colored individual trajectories overlapped frame by frame. Scale bar: 500µm

**Supplementary Movie M3**. Two-hour movie of the random motility experiment presented in Fig. 2 a (cropped FOV), with time-normalized individual trajectories overlapped frame by frame. Microscope: lens-free Cytonote 6W, IPRASENSE with 1.68 µm/pixel resolution and a time interval of 45s between consecutive frames. Scale bar: 500 µm

## References

1. A. D. Luster, R. Alon, U. H. von Andrian, Immune cell migration in inflammation: present and future therapeutic targets. Nat. Immunol. 6, 1182–1190 (2005).

2. P. Friedl, D. Gilmour, Collective cell migration in morphogenesis, regeneration and cancer. Nat. Rev. Mol. Cell Biol. 10, 445–457 (2009).

3. E. S. Gloag, et al., Self-organization of bacterial biofilms is facilitated by extracellular DNA. Proc. Natl. Acad. Sci. 110, 11541–11546 (2013).

4. R. H. Kessin, Dictyostelium: Evolution, Cell Biology, and the Development of Multicellularity, C. U. Press, Ed. (2001).

5. F. W. Rossine, G. T. Vercelli, C. E. Tarnita, T. Gregor, Structured foraging of soil predators unveils functional responses to bacterial defenses. Proc. Natl. Acad. Sci. U. S. A. 119, e2210995119 (2022).

6. C. M. Waters, B. L. Bassler, Quorum sensing: cell-to-cell communication in bacteria. Annu. Rev. Cell Dev. Biol. 21, 319–346 (2005).

7. J. Hickson, et al., Societal interactions in ovarian cancer metastasis: a quorum-sensing hypothesis. Clin. Exp. Metastasis 26, 67–76 (2009).

8. H. Jayatilaka, et al., Synergistic IL-6 and IL-8 paracrine signalling pathway infers a strategy to inhibit tumour cell migration. Nat. Commun. 8, 15584 (2017).

9. T. Gregor, The Onset of Collective Behavior. 1021 (2011).

10. S. S. Willard, P. N. Devreotes, Signaling pathways mediating chemotaxis in the social amoeba, Dictyostelium discoideum. Eur. J. Cell Biol. 85, 897–904 (2006).

11. L. Golé, C. Rivière, Y. Hayakawa, J. P. Rieu, A quorum-sensing factor in vegetative dictyostelium discoideum cells revealed by quantitative migration analysis. PLoS One 6, e26901 (2011).

12. J. D’Alessandro, et al., Collective regulation of cell motility using an accurate density-sensing system. J. R. Soc. Interface 15, 20180006 (2018).

13. J. D’alessandro, et al., Contact enhancement of locomotion in spreading cell colonies. Nat. Phys. 13, 999–1005 (2017).

14. M. Abercrombie, J. E. M. Heaysman, Observations on the social behaviour of cells in tissue culture. Exp. Cell Res. 6, 293–306 (1954).

15. R. Mayor, C. Carmona-Fontaine, Keeping in touch with contact inhibition of locomotion. Trends Cell Biol. 20, 319–328 (2010).

16. T. Fujimori, A. Nakajima, N. Shimada, S. Sawai, Tissue self-organization based on collective cell migration by contact activation of locomotion and chemotaxis. Proc Natl Acad Sci U S A 2, 1–25.

17. A. Shellard, R. Mayor, Rules of collective migration: from the wildebeest to the neural crest. Phil. Trans. R. Soc. B 375, 20190387 (2020).

18. T. E. Angelini, E. Hannezo, X. Trepat, J. J. Fredberg, D. A. Weitz, Cell Migration Driven by Cooperative Substrate Deformation Patterns. Science (80-.). 168104, 1–4 (2010).

19. S. Vedel, S. Tay, D. M. Johnston, H. Bruus, S. R. Quake, Migration of cells in a social context. Proc. Natl. Acad. Sci. U. S. A. 110, 129–134 (2013).

20. N. Guisoni, K. I. Mazzitello, L. Diambra, Modeling Active Cell Movement With the Potts Model. Front. Phys. 6 (2018).

21. O. Cochet-Escartin, et al., Hypoxia triggers collective aerotactic migration in Dictyostelium discoideum. Elife 10, e64731 (2021).

22. R. Siddiqui, R. Dudley, N. A. Khan, Acanthamoeba differentiation: a two-faced drama of Dr Jekyll and Mr Hyde. Parasitology 139, 826–834 (2012).

23. N. A. Kuburich, N. Adhikari, J. A. Hadwiger, Acanthamoeba and Dictyostelium Use Different Foraging Strategies. Protist 167, 511–525 (2016).

24. F. L. Schuster, M. Levandowsky, Chemosensory Responses of Acanthamoeba castellanii: Visual Analysis of Random Movement and Responses to Chemical Signals. J. Eukaryot. Microbiol. 43, 150–158 (1996).

25. R. J. Neff, Purification, Axenic Cultivation, and Description of a Soil Amoeba, Acanthamoeba sp. J. Protozool. 4, 176–182 (1957).

26. S. Berg, et al., ilastik: interactive machine learning for (bio)image analysis. Nat. Methods 16, 1226–1232 (2019).

27. D. Allan, T. Caswell, N. Keim, C. van der Wel, R. Verweij, soft-matter/trackpy: v0.6.4 (v0.6.4) (Zenodo. 10.5281/zenodo.12708864, 2024).

28. C. R. Harris, et al., Array programming with NumPy. Nature 585, 357–362 (2020).

29. W. McKinney, {D}ata {S}tructures for {S}tatistical {C}omputing in {P}ython in {P}roceedings of the 9th {P}ython in {S}cience {C}onference, S. van der Walt, J. Millman, Eds. (2010), pp. 56–61.

30. J. D. Hunter, Matplotlib: A 2D Graphics Environment. Comput. Sci. Eng. 9, 90–95 (2007).

31. P. Virtanen, et al., SciPy 1.0: fundamental algorithms for scientific computing in Python. Nat. Methods 17, 261–272 (2020).

32. J. Schindelin, et al., Fiji: an open-source platform for biological-image analysis. Nat. Methods 9, 676–682 (2012).

33. R. B. Dickinson, R. T. Tranquillo, Optimal estimation of cell movement indices from the statistical analysis of cell tracking data. AIChE J. 39, 1995–2010 (1993).

34. J. M. Ver Hoef, Who Invented the Delta Method? Am. Stat. 66, 124–127 (2012).

35. J. P. Rieu, A. Upadhyaya, J. A. Glazier, N. B. Ouchi, Y. Sawada, Diffusion and Deformations of Single Hydra Cells in Cellular Aggregates. Biophys. J. 79, 1903–1914 (2000).

36. E. A. Codling, M. J. Plank, S. Benhamou, Random walk models in biology. Interface, 813–834 (2008).

37. G. E. Uhlenbeck, L. S. Ornstein, On the Theory of the Brownian Motion. Phys. Rev. 36, 823–841 (1930).

38. G. L. Thomas, et al., Parameterizing cell movement when the instantaneous cell migration velocity is ill-defined. Phys. A Stat. Mech. its Appl. 550, 124493 (2020).

39. N. Ghazi, “Aerotaxis and the collective spreading of social and asocial amoebae,” Université Claude Bernard Lyon1. (2024).

40. R. P. Feynman, R. B. Leighton, M. Sands, The Feynman lectures on physics; New millennium ed. (Basic Books, 2010).

41. C. Bechinger, et al., Active particles in complex and crowded environments. Rev. Mod. Phys. 88, 45006 (2016).

42. A. Dehkharghani, N. Waisbord, J. S. Guasto, Self-transport of swimming bacteria is impaired by porous microstructure. Commun. Phys. 6, 18 (2023).

43. D. Arcizet, et al., Contact-controlled amoeboid motility induces dynamic cell trapping in 3D-microstructured surfaces. Soft Matter 8, 1473 (2012).

44. S. Hirose, J. Rieu, O. Cochet-escartin, C. Anjard, K. Funamoto, The Oxygen Gradient in Hypoxic Conditions Enhances and Guides Dictyostelium discoideum Migration. Processes 10, 318 (2022).

